# A Role for Cross-linking Proteins in Actin Filament Network Organization and Force Generation

**DOI:** 10.1101/2024.04.19.590161

**Authors:** Jennifer M. Hill, Songlin Cai, Michael D. Carver, David G. Drubin

## Abstract

The high turgor pressure across the plasma membrane of yeasts creates a requirement for substantial force production by actin polymerization and myosin motor activity for clathrin-mediated endocytosis (CME). Endocytic internalization is severely impeded in the absence of fimbrin, an actin filament crosslinking protein called Sac6 in budding yeast. Here, we combine live-cell imaging and mathematical modeling to gain new insights into the role of actin filament crosslinking proteins in force generation. Genetic manipulation showed that CME sites with more crosslinking proteins are more effective at internalization under high load. Simulations of an experimentally constrained, agent-based mathematical model recapitulate the result that endocytic networks with more double-bound fimbrin molecules internalize the plasma membrane against elevated turgor pressure more effectively. Networks with large numbers of crosslinks also have more growing actin filament barbed ends at the plasma membrane, where the addition of new actin monomers contributes to force generation and vesicle internalization. Our results provide a richer understanding of the crucial role played by actin filament crosslinking proteins during actin network force generation, highlighting the contribution of these proteins to the self-organization of the actin filament network and force generation under increased load.

## Introduction

Clathrin-mediated endocytosis (CME) is the process by which cells take up membrane-bound and soluble molecules from their surface and environment through the internalization of a clathrin-coated vesicle. CME is highly conserved from budding yeast to mammalian cells, as it is a major pathway for internalizing necessary cellular building blocks and down-regulating signal transduction by membrane receptors. The process can be described as consisting of a set of modules, each composed of proteins that work in concert to recruit cargo, adaptors, and coat proteins, build an actin filament network to invaginate the membrane against resistive membrane forces, and release the nascent vesicle into the cytoplasm (1, 2). In budding yeast, the proteins in each of these modules and their biochemical properties are well-characterized, which, in addition to the high level of conservation to mammalian CME, makes it an ideal organism for studying this process (3).

While actin filament assembly has been associated with CME from yeast to humans (4, 5), the high turgor pressure in yeast creates a particularly pronounced requirement for forces generated by polymerization of a branched actin network and myosin motor activity (6–8). The Arp2/3 complex nucleates a branched actin network when activated by the yeast WASP, also known as Las17, and the CA domain of type-I myosins (7, 9). Adaptor proteins Sla2 and Ent1/2 couple the actin filament network to the growing membrane invagination (10). Additional actin-associated proteins such as myosins (7, 11), capping proteins (12), the actin-filament severing protein cofilin (13), and actin crosslinking proteins (2, 14) all play important roles in the process, but how these proteins work together to build an actin network capable of force production to drive membrane bending is still not fully understood.

At least two actin crosslinking proteins are involved in CME in budding yeast. The yeast fimbrin, Sac6, and the yeast transgelin, Scp1, crosslink actin filaments *in vitro*, and are localized to the actin filament network at endocytic sites (15, 16). While fimbrin and transgelin undergo genetic dosage compensation (17), the effect of their relative expression levels on their recruitment to and function at endocytic sites has not been reported. Yeast fimbrin is essential for CME (1), and its absence leads to a growth defect (16). Additionally, actin networks assembled *in vitro* from yeast extracts lacking fimbrin have a lower elastic modulus than networks assembled in wild-type yeast extracts, indicating that fimbrin contributes to the stiffness of actin networks *in vitro* (18, 19). The absence of transgelin exacerbates the endocytic defects observed in cells lacking fimbrin, but the absence of transgelin alone has only a minor effect on cell growth and endocytosis (16) or the elastic modulus of actin filament networks assembled in yeast extracts (18). For these reasons, transgelin is a relatively under- studied component of budding yeast CME machinery.

While CME is a highly robust process, actin filament crosslinking is essential for the formation of actin networks capable of carrying out this complex membrane bending process (1, 4). Crosslinking proteins participate in most actin-mediated processes, but how actin filament crosslinking affects the geometry and force-generating capabilities of actin networks is still not fully understood. The role of crosslinking proteins in CME has been particularly difficult to study because of the dynamic nature of the process and because CME sites are diffraction-limited, meaning that details of actin filament organization cannot be resolved using conventional or even super-resolution light microscopy techniques. From yeast to humans, the actin filament network at CME sites assembles and disassembles on the time scale of ten to 20 seconds (1, 4). Electron microscopy has been used to visualize the geometry of the endocytic actin network in mammalian cells (20, 21) but lacks information about the dynamics of the process.

Given these limitations, mathematical models are highly attractive tools for predicting novel functions of actin crosslinking proteins. When combined with *in vivo* experiments, they can provide deeper mechanistic insights than either technique used alone.

In this study, we investigated the role of actin filament crosslinking proteins in clathrin-mediated endocytosis in budding yeast, focusing on their effects on actin filament organization and force generation. We used quantitative microscopy to determine the numbers of yeast fimbrin and yeast transgelin molecules at CME sites in cells grown under various conditions and tested for an association between the number of crosslinking proteins and endocytic success at high turgor pressure. Markedly more fimbrin is present at CME sites than transgelin, and overexpression of transgelin can at least partially rescue CME in cells lacking fimbrin. Additional fimbrin can assist in internalization during the adaptive response to increased turgor pressure resulting from hypotonic shock, revealing a role for crosslinking proteins in force generation under high load during CME. Simulations generated by a mathematical model of the actin filament network during CME in yeast showed that crosslinking proteins promote actin filament network self-organization and concentrate growing filament ends at the plasma membrane, producing higher forces and more efficient internalization. These studies identified novel roles for actin filament crosslinking proteins in force production and actin network organization during budding yeast CME.

## Results

### Interplay between fimbrin and transgelin actin filament crosslinking protein recruitment to CME sites

The maximum number of molecules of the most abundant actin filament crosslinking protein at CME sites, fimbrin, has previously been estimated using quantitative fluorescence microscopy (22). We used the same quantitative microscopy method to measure the maximum number of transgelin molecules arriving at endocytic sites. Transgelin is a less abundant actin filament crosslinking protein. We used the ratio between the maximum fluorescence intensities of endogenously labeled Las17- GFP, a protein whose abundance at endocytic sites has been measured (22), and GFP-tagged transgelin in the same field of view to determine that the maximum number of transgelin molecules recruited to CME sites is 40±15 (Fig. S1A, S1B).

Transgelin protein levels have been reported to increase upon deletion of the gene encoding yeast fimbrin, *SAC6* (17). We tested how fimbrin absence affects the number of transgelin molecules at CME sites and found that more transgelin molecules are recruited to CME sites in cells lacking fimbrin (58±40, p<0.0005) relative to CME sites in wild-type cells (Fig. 1A, 1B). This observation suggests that dosage compensation by transgelin for fimbrin absence contributes to increased transgelin recruitment to CME sites. We also found that transgelin lifetime at CME sites increases in the absence of fimbrin, consistent with previously published results showing that absence of fimbrin leads to increased actin patch lifetimes (1, 17) (Fig. S1C).

**Fig. 1.**
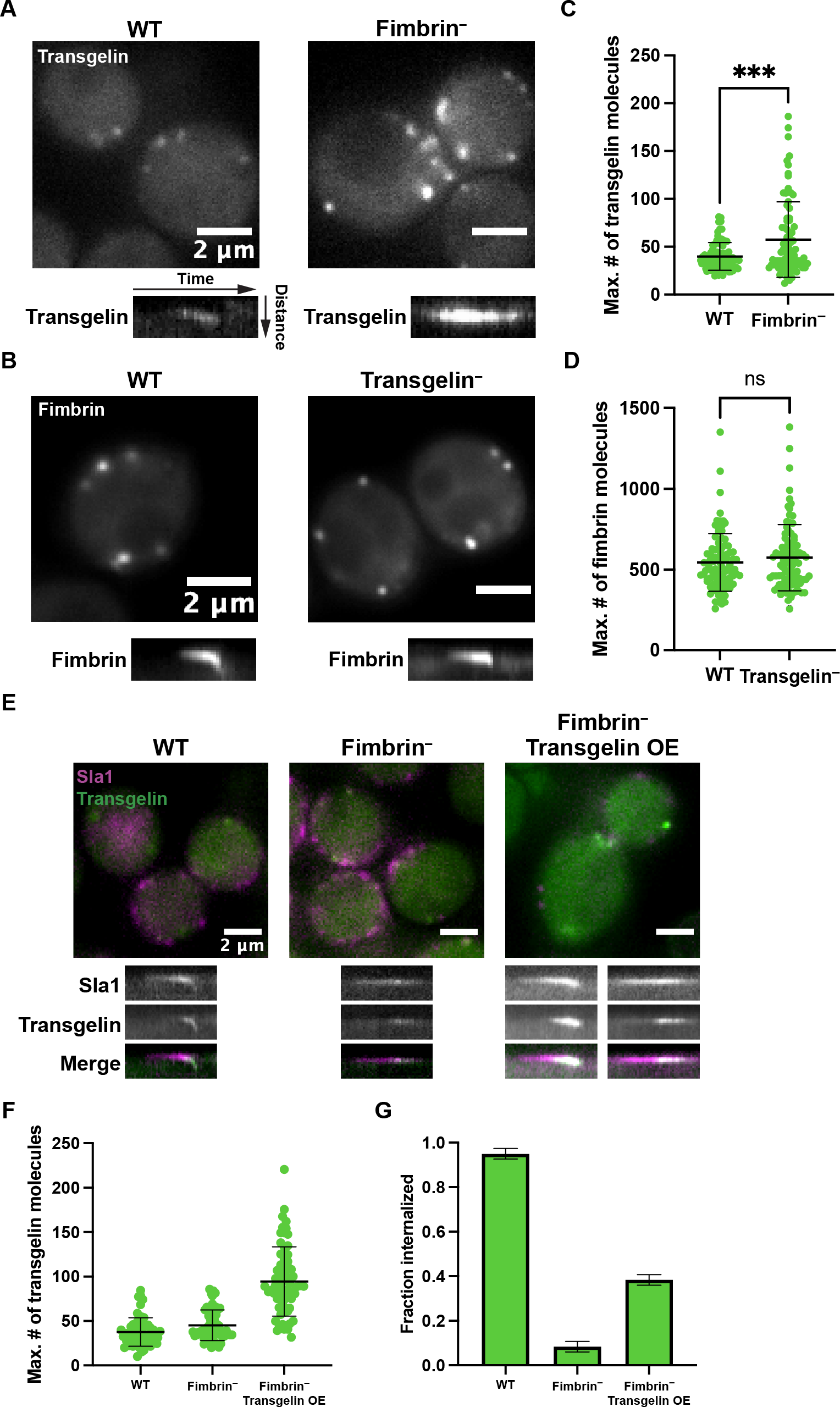
Loss of fimbrin leads to an increase in the number of molecules of the minor actin filament crosslinking protein transgelin, but loss of transgelin does not affect the number of fimbrin molecules at CME sites. (A), (B). Quantitative microscopy to determine the maximum number of GFP-tagged transgelin (A) or fimbrin (B) molecules at sites of CME in wild-type strains and strains lacking fimbrin (fimbrin^−^) or transgelin (transgelin^−^), respectively. Kymographs of individual sites show a slice from outside the cell (top) to the inside (bottom) over time. (C), (D). Maximum number of molecules of transgelin (C) and fimbrin (D) at CME sites in wild-type and fimbrin^−^ or transgelin^−^ strains, respectively (n=90 sites from ≥10 cells per strain). (E). Representative images of mCherry-tagged Sla1 and GFP-tagged transgelin in WT, fimbrin^−^, and fimbrin^−^ transgelin overexpression (transgelin OE) strains, 24 hours post-induction with β- estradiol. Sites where Sla1 and transgelin hook into the cell in kymographs were considered internalized. Representative kymographs are shown for all strains. (F). Quantification of maximum numbers of transgelin molecules arriving at CME sites in WT, fimbrin^−^, and fimbrin^−^ transgelin OE strains, 24 hours post-induction with β-estradiol (n=60 sites from ≥10 cells per strain). (G). Fraction of CME sites internalized in WT, fimbrin^−^, and fimbrin^−^ transgelin OE strains, 24 hours post-induction with β-estradiol (n=60 sites from ≥10 cells per strain). Error bars show standard deviation.

Next, we investigated whether previously reported increased fimbrin expression when transgelin is absent similarly increases fimbrin recruitment to CME sites. In wild-type cells, the maximum number of fimbrin molecules at CME sites was previously determined to be 545±135 (22). Using the fluorescence intensity of GFP-tagged fimbrin in wild-type cells as the standard, the number of fimbrin molecules recruited to endocytic sites in cells lacking transgelin was slightly higher than in wild-type cells, although the difference was not statistically significant (574±205 p=0.3087). Thus, in the absence of transgelin, changes in fimbrin recruitment to CME sites were at best small (Fig. 1C, 1D). Additionally, transgelin absence did not significantly increase the fimbrin lifetime at CME sites, consistent with a previously reported modest endocytic defect observed in cells lacking transgelin (16) (Fig. S1D). These results indicate that the reduced requirement for transgelin in crosslinking actin filaments at CME sites relative to fimbrin might be at least partially explained by its relatively low abundance at CME sites compared to fimbrin.

We then asked whether overexpression of transgelin can rescue the endocytic defect caused by the absence of fimbrin. We overexpressed GFP-tagged transgelin in the absence of fimbrin and quantified the number of transgelin molecules arriving at CME sites. Incorporation of the *GAL1* promoter upstream of the gene encoding transgelin, *SCP1*, disrupted the native promoter, resulting in negligible expression of transgelin before induction of overexpression (Fig S1E). However, induction of transgelin overexpression results in the recruitment of 94±39 molecules of transgelin to CME sites, ∼2.4 times the recruitment of transgelin in wild-type cells and cells lacking fimbrin (Fig. 1E, 1F). We used the endocytic coat marker, Sla1, to measure the fraction of CME sites that successfully internalize in these conditions. Wild-type cells internalize ∼95±2% of CME sites, while cells lacking fimbrin internalize only ∼8±2% of sites, indicating a severe endocytic defect (Fig. 1E, Fig. 1G, and Movie S1). However, cells overexpressing transgelin internalize ∼38±2% of CME sites (Fig. 1E, Fig. 1G, and Movie S1), indicating that increased recruitment of transgelin to CME sites can partially rescue the endocytic defect caused by the absence of fimbrin. Taken together, these results show that the relative abundance of crosslinking proteins at CME sites, as opposed to the identity of the crosslinking proteins, is an important factor in the success of CME.

### Mathematical modeling reveals a force-generating role for actin filament crosslinking proteins

Compensatory recruitment of more transgelin to CME sites in cells lacking fimbrin suggests that the number of crosslinking proteins recruited to CME sites might be important for endocytic success. To investigate this possibility, we adapted an agent-based mathematical model of the actin network during yeast CME (9) to simulate the effects of a range of numbers of crosslinking proteins on vesicle internalization. The model approximates the internalizing vesicle as a cylinder with a hemispherical end attached to a spring that provides a force resistive to internalization (Fig. 2A). The magnitude of this force was set based on models of membrane mechanics (23) such that the actin filament network must overcome an initial force barrier, followed by a linear increase in the force requirement proportional to the distance the vesicle has internalized (9). Actin filaments are attached to the tip of the vesicle by a strong actin linker and grow radially outward from this point (Fig. 2A). Pre-activated Arp2/3 complex diffuses at the membrane and binds to the growing actin filaments to nucleate daughter filaments at a branching angle of 70° (Fig. 2A). Crosslinking proteins diffuse in the cytosolic space and can bind to two actin filaments to form a crosslink (Fig. 2A). Both the Arp2/3 complex and crosslinking proteins stochastically bind and unbind actin filaments at rates determined experimentally in *in vitro* studies of the Arp2/3 complex and fimbrin (9).

**Fig. 2.**
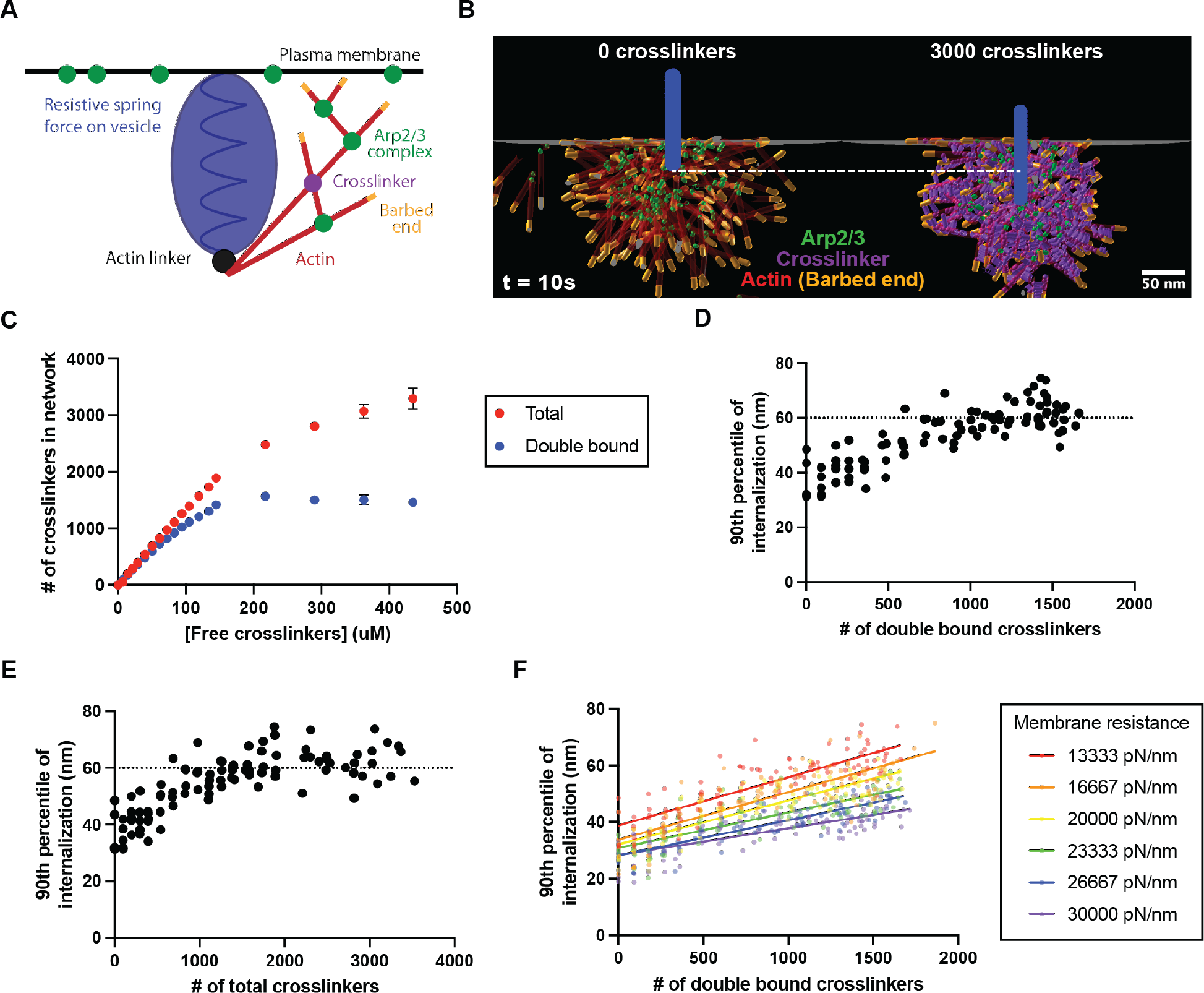
Mathematical modeling reveals that crosslinking proteins contribute to actin network force generation for membrane internalization during CME. (A) Schematic of an agent-based model for the actin network associated with CME sites (9) The endocytic vesicle is modeled as a bead on a spring (blue) that resists internalization. Actin filaments attached to the vesicle generate force for internalization by polymerizing at the membrane. (B) Final frame of simulations with pools of either 0 or 3000 available crosslinking proteins. The view shows a slice from the center of the network to reveal the internalized vesicle in the network (blue). (C) Numbers of total and double-bound crosslinkers in the final actin network across a range of free crosslinker concentrations. N=5 simulations, error bars show standard deviation. (D) Scatter plot of the 90^th^ percentile of the extent of vesicle internalization for simulations with a range of numbers of double-bound crosslinkers in the final network. Dotted line at Y=60 nm represents threshold for snap-through transition and scission. (E) Scatter plot of the 90^th^ percentile of the extent of vesicle internalization for simulations with varying numbers of total crosslinkers in the final network. Dotted line at Y=60 nm represents threshold for snap- through transition and scission. (F) Scatter plot of the 90^th^ percentile of the extent of internalization for individual simulations with varying numbers of double-bound crosslinkers and membrane resistances. Solid lines show linear regressions for each set of simulations at a given membrane resistance.

In the absence of crosslinking proteins, actin filaments that grow toward the plasma membrane encounter and bind to the Arp2/3 complex, which nucleates daughter filaments, generating an expanding network of branched actin filaments that generates force to internalize the vesicle throughout a ten-second simulation (Fig. 2B, left, and movie S2). We observed that in the presence of crosslinking proteins, crosslinks are formed between actin filaments, and the branched actin filament network internalizes the vesicle further (see white dotted reference line) than in the absence of crosslinking proteins (Fig. 2B, right, and movie S2). In all simulations without crosslinking proteins, actin filaments dissociated from the actin network due to Arp2/3 unbinding from the mother filament, while this was never observed in simulations with a pool of 3000 available crosslinking proteins (Fig. 2B).

To investigate the effect of the density of crosslinks on the endocytic actin network, we ran simulations with a range of concentrations of crosslinking proteins in the simulation space. The number of crosslinking proteins bound to at least one actin filament in the network after ten seconds increases with the concentration of free crosslinking proteins (Fig. 2C, S2A), but the number of crosslinking proteins binding to two actin filaments to form a crosslink (double-bound crosslinkers), plateaus at around 2000 total crosslinking proteins (Fig. 2C, S2B). These data suggest that there is an upper limit to the number of double-bound crosslinking proteins that can exist in the branched actin filament network.

We then investigated the effect of the density of double-bound crosslinking proteins on the extent of invagination. To reduce stochastic noise in vesicle internalization, we calculated the 90th percentile of internalization over all simulated time points rather than using the internalized distance at the final time point of the simulation. The extent of invagination increased in proportion to the number of double-bound crosslinkers (Fig. 2D, S2C), suggesting that crosslinking of actin filaments within the network enhances force generation during CME internalization. Modeling of membrane mechanics suggests that an energetic snap-through transition occurs when the invagination reaches 60 nm, at which point scission occurs (24), and the majority of simulations with 1300-1500 double-bound crosslinkers reached this point (Fig. 2D). To ensure that this effect is specifically due to the formation of actin filament crosslinks, rather than simply the result of more crosslinking proteins binding in the actin filament network, we also measured the effect of total bound crosslinking proteins in the final network on vesicle internalization. As the total number of bound crosslinking proteins increases to around 2000, internalization of the vesicle also increases, corresponding to the range over which double-bound crosslinkers are increasing in the network (Fig. 2E). However, increasing the total number of crosslinking proteins beyond 2000 had little to no effect on the internalization of the invagination, indicating that crosslinking proteins must form filament crosslinks to enhance force generation for internalization.

Interestingly, the effect of crosslinking proteins on internalization is similar across a wide range of force regimes. By increasing the spring constant used to simulate the plasma membrane’s resistance to internalization, we modeled the effect of increasing turgor pressure or membrane tension. Networks generated under increased membrane tension conditions still recruit similar numbers of total and double-bound crosslinkers as in simulations with lower membrane tension, plateauing at around 1400-1500 double-bound crosslinkers (Fig. S2D, S2E), and these networks do not internalize the membrane as far as those generated with lower membrane tension (Fig. 2F). However, at each force regime, internalization still increases with the density of double-bound crosslinkers in the network (Fig. 2F, S2F). Taken together, these results suggest that increasing the number of double-bound crosslinkers increases force generation and internalization over a wide range of resistive forces.

### Actin filament crosslinking protein importance for CME internalization tested under high turgor pressure *in vivo*

To test the model prediction that actin filament crosslinking proteins contribute to force generation even under conditions of high resistive forces, we next investigated the effect of varying numbers of crosslinking proteins on the efficiency of CME *in vivo*. In the following experiments, we deleted the gene encoding transgelin to eliminate the complication of differing contributions from multiple types of crosslinking proteins. We used hypotonic shock to increase turgor pressure to the point when about half of the CME events failed (25, 26), and compared numbers of fimbrin molecules at sites that successfully internalize or fail to internalize. We also deleted the gene encoding the glycerol export protein, Fps1, which is important to restore membrane tension in response to hypotonic shock, extending the time during which cells have increased turgor pressure (25, 27). Before shock, the fraction of endocytic sites that internalize in wild-type cells, using the endocytic coat protein Sla1 as a marker for internalization, is 0.89±0.07, while after hypotonic shock, the fraction of sites that internalize is 0.53±0.14 (Fig. S3A, S3B). The lifetimes of Sla1 and the actin-binding protein Abp1 at CME sites also increase upon hypotonic shock, indicative of a defect in CME (Fig. S3C, S3D).

We then generated strains with varying copy numbers of the fimbrin gene, *SAC6*: a heterozygous diploid (*SAC6/sac6Δ*), a wild-type diploid (*SAC6/SAC6*), a wild-type haploid (*SAC6*), and a haploid bearing a duplication of the gene encoding fimbrin at the endogenous locus (*SAC6* dupl.). We used quantitative microscopy to measure the maximum number of fimbrin molecules arriving at endocytic sites after hypotonic shock in these strains and found a range of numbers of fimbrin molecules at each site (Fig. 3A, 3C, and S3E). We also quantified the fraction of endocytic sites that internalize in each of these strains after hypotonic shock and found that strains with an increased copy number of the gene encoding fimbrin have a higher fraction of sites that internalize compared to strains with a lower gene dosage (Fig. 3B, 3C, S3F, and Movie S3). Taken together, these data corroborate the model prediction that increased numbers of crosslinking proteins contribute to internalization under conditions of high turgor pressure and suggest that crosslinking proteins may be involved in an adaptive response to provide additional force for internalization.

**Fig. 3.**
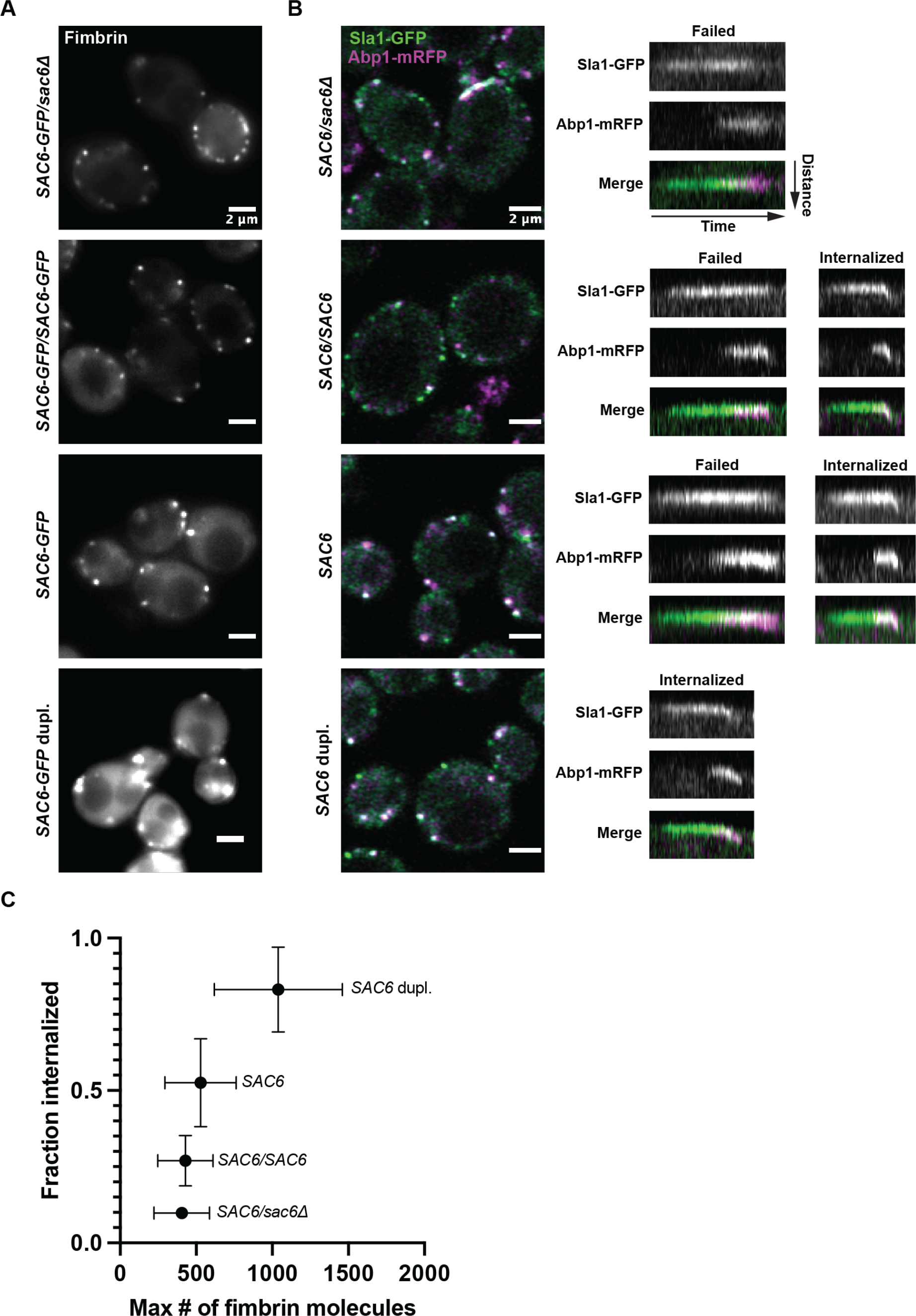
The extent to which CME site internalization is hindered by hypotonic shock depends on crosslinking protein abundance. Cells were grown in media containing 1M sorbitol and shifted to media containing 0.25M sorbitol for five minutes. All strains are in a background containing a deletion of the genes encoding transgelin and Fps1. *SAC6* dupl.: Duplication of the gene encoding fimbrin at the endogenous locus. (A) Quantitative imaging of GFP-tagged fimbrin five minutes after osmotic shock. (B) Imaging of GFP-tagged Sla1 and mRFP-tagged Abp1 five minutes after osmotic shock. Sites where Sla1 and Abp1 hook into the cell in kymographs were considered internalized. Representative kymographs are shown for all strains. (C) Fraction of endocytic sites that are internalized as a function of the maximum number of fimbrin molecules at endocytic sites five minutes post-shock (n=90 sites from ≥10 cells per strain). Error bars show standard deviation.

To investigate the relationship between crosslinking proteins and CME efficiency at individual CME sites, we categorized endocytic events as internalized or failed based on the movement of both fimbrin and Abp1 away from the membrane and quantified the maximum number of fimbrin molecules recruited to these sites (Fig. 4A and Movie S4). Sites that internalize have significantly more crosslinking proteins than sites that fail to internalize (Fig. 4B). The same result was observed using Sla1, associated with the vesicle coat rather than an actin network component, as a marker for endocytic internalization (data not shown). Additionally, we also observed that sites that successfully internalize have significantly higher Abp1 fluorescence intensity, likely reflecting more assembled actin, than sites that fail to internalize (Fig. 4C). However, the ratio of the number of fimbrin molecules to Abp1 fluorescence intensity is higher for sites that internalize, indicating that these actin filaments are more saturated with crosslinking proteins than actin filaments at sites that fail to internalize (Fig. 4D). Taken together, these data demonstrate that CME sites that recruit more crosslinking proteins are better able to internalize and show that the importance of actin filament crosslinking proteins for successful CME internalization is increased under conditions of high turgor pressure.

**Fig. 4.**
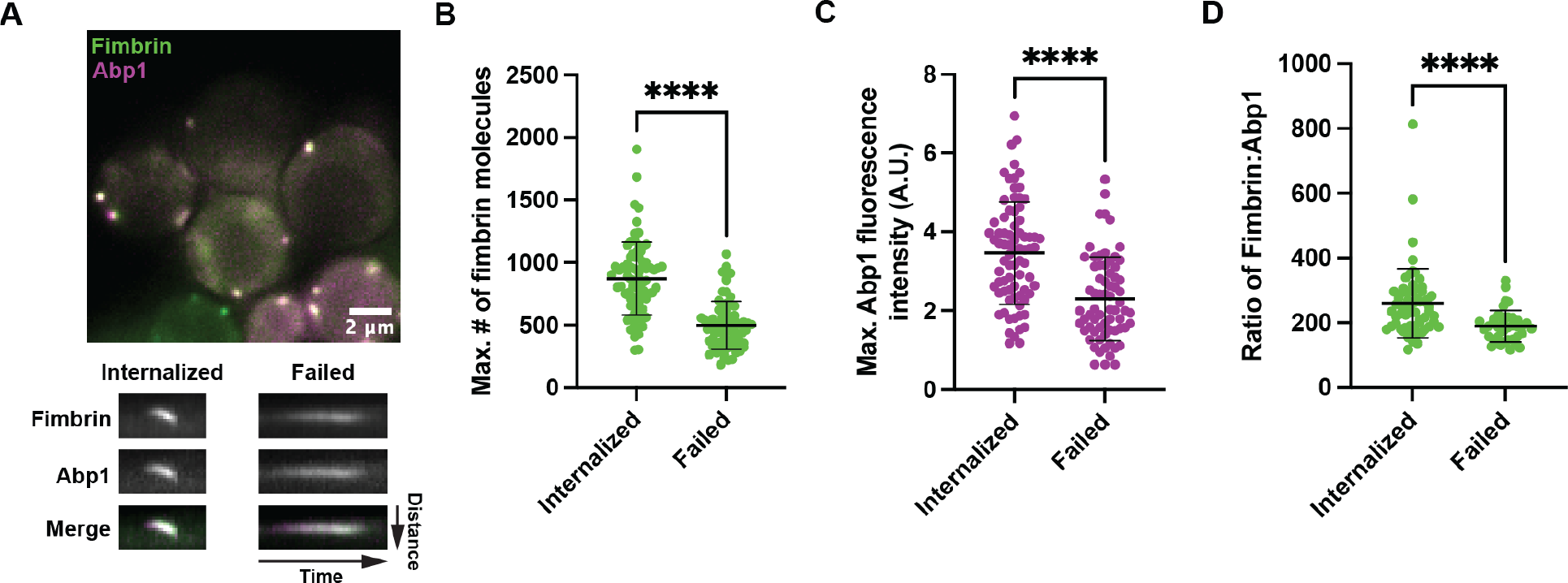
Endocytic sites that internalize under hypotonic shock conditions have a higher maximum number of fimbrin molecules than sites that stall. (A) Cells were grown in 1M sorbitol media and shifted to 0.25M sorbitol media. Fimbrin and Abp1 were imaged after five minutes. (B) Maximum number of fimbrin molecules at endocytic sites that internalize (n=62 sites) or fail (n=38 sites). (C) Maximum fluorescence intensity of mRFP-tagged Abp1 at endocytic sites that internalize (n=62 sites) or fail (n=38 sites). (D) Ratio of the maximum number of fimbrin molecules to Abp1-mRFP fluorescence intensity at endocytic sites that internalize (n=62 sites) or fail (n=38 sites). Error bars show standard deviation.

### Actin filament crosslinking proteins organize the actin network for maximal force production

To investigate how crosslinking proteins contribute to force generation by the actin filament network during CME, we analyzed the organization of actin filaments in simulations with different numbers of crosslinking proteins. Since growing actin filaments can generate forces for internalization, we defined a growing end enrichment score, which represents the enrichment of growing ends in a volume of the actin network at the end of the ten-second simulation. The metric was calculated by subtracting the number of capped actin filament ends in a volume of the network from the number of growing actin filament ends in the same volume. A positive growing end enrichment score indicates a region of the network with more growing than capped actin filament ends. Without crosslinking proteins, the growing end enrichment score is negative for all network regions (Fig. 5A, left), indicating that there are no regions of these networks that are enriched for growing ends. However, in simulations starting with 3000 free crosslinkers, equivalent to about 1600 double-bound crosslinkers in the final actin network, the growing end enrichment score is positive in the region of the actin network where filaments contact the membrane (Fig. 5A, right), leading to an enrichment of growing filaments ends at the membrane where they can generate force for internalization. This enrichment effect becomes more pronounced throughout the invagination process, such that the final actin network has the highest enrichment of growing ends at the membrane when the membrane is invaginated the most (Fig. S4A). Increasing the number of crosslinking proteins also enriches the number of growing ends at the membrane under high membrane tension conditions (Fig. 5B). Interestingly, we found that the number of crosslinking proteins has very little effect on the overall axial orientation of actin filaments in the network with respect to the flat membrane, meaning that crosslinking of the actin network does not significantly reorient actin filaments in any direction (Fig. S4B, S4C). This result indicates that crosslinking proteins assist in the formation of an actin filament network that is self-organized to generate force through the enrichment of growing filament ends at the plasma membrane.

**Fig. 5.**
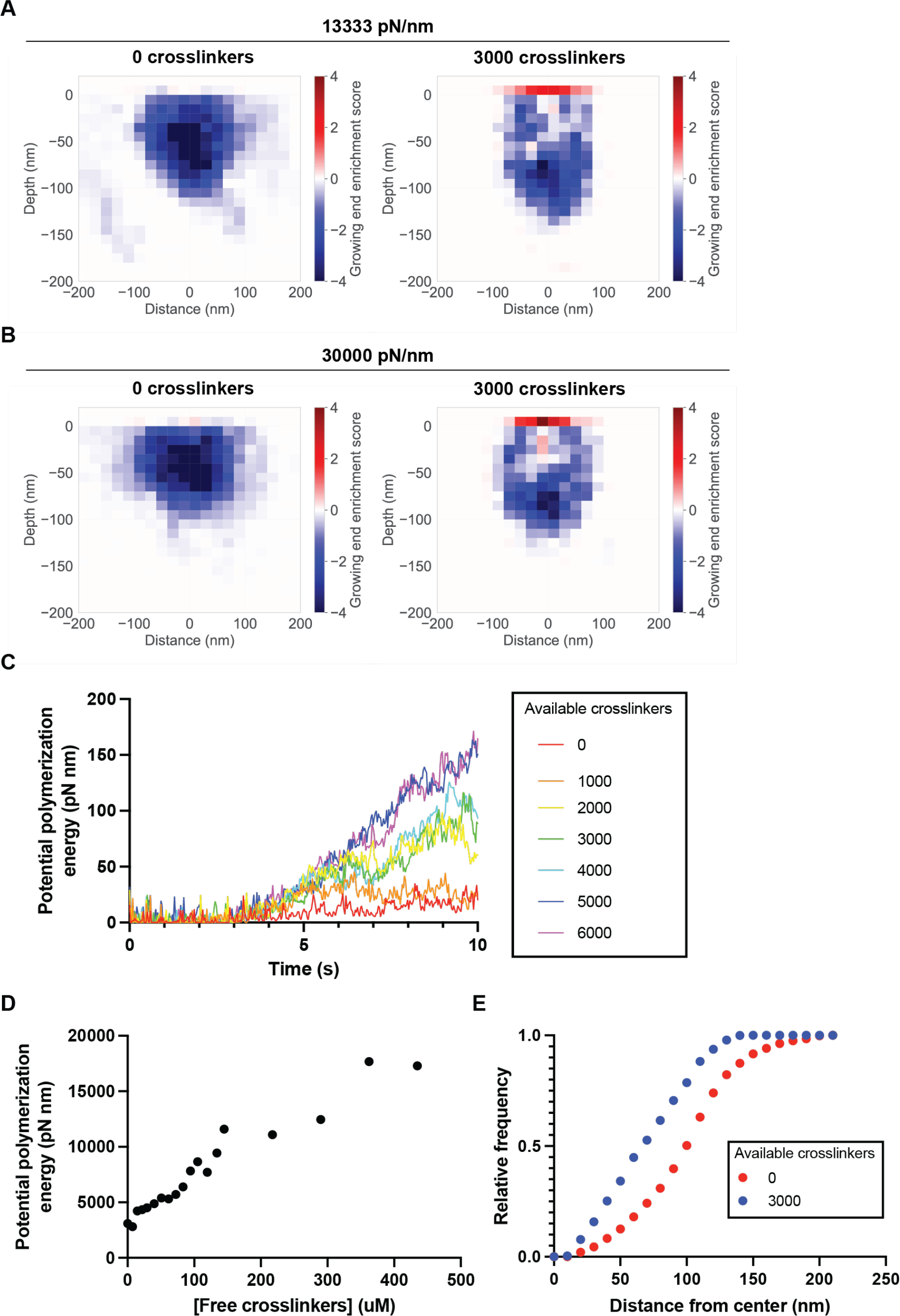
Mathematical modeling predicts self-organization of actin filaments to maximize force production. (A-B) Heat map of enrichment scores for growing ends relative to capped ends of actin filaments in the final network with pools of either 0 or 3000 available crosslinking proteins and 13333 pN/nm (A) or 30000 pN/nm (B) required to internalize the vesicle. N=5 simulations. (C) Average potential polymerization energy of the actin filament network over the duration of simulations with varying pools of available crosslinking proteins. N=5 simulations. (D) Total potential polymerization energy of the actin filament network as a function of the concentration of free crosslinkers. N=5 simulations. (E) Cumulative distribution functions for the relative frequency of plus ends measured radially from the center of the vesicle for simulations with pools of 0 and 3000 available crosslinking proteins. N=5 simulations.

Growing end enrichment at the plasma membrane in response to elevated numbers of crosslinking proteins suggests that with more crosslinking proteins, the actin filament network has a higher potential energy to be harnessed for force generation. We defined potential polymerization energy as the sum of the polymerization energies of each growing actin filament in contact with the membrane at any given time. Over time, as the network grows and more filaments encounter the membrane, the potential polymerization energy increases (Fig. 5C). However, simulations with a larger pool of available crosslinking proteins generate networks that increase their potential polymerization energy at a faster rate than is observed in simulations with fewer crosslinking proteins (Fig. 5C). Interestingly, in simulations with small pools of available crosslinking proteins, the potential polymerization energy of the actin filament network plateaus as crosslinking proteins become saturated in the network (Fig. 5C, S4D, and S4E). Additionally, we defined the total polymerization energy of the network as the sum of the potential polymerization energies at each time point. As the free crosslinking protein concentration increases, the total polymerization energy of the network throughout its lifetime increases (Fig. 5D), suggesting that crosslinking proteins contribute to the generation of a network with higher potential energy to be harnessed upon actin polymerization.

We also observed the overall compaction of the actin network in simulations with more double-bound crosslinkers, specifically at the contact area between the actin network and the membrane. By quantifying the positions of growing filament ends at the membrane measured radially from the center of the internalizing vesicle in the final second of the simulation, we found that increasing crosslinking protein levels decreases the contact area of filament ends at the plasma membrane such that filament ends are localized closer to the center of the internalizing membrane tubule (Fig. 5E). Taken together, these data reveal a role for actin crosslinking proteins in the organization of a compacted actin network enriched in growing filament ends at the membrane, increasing the probability that actin polymerization leads to successful vesicle internalization.

## Discussion

In this study, we set out to determine the role of actin filament crosslinking proteins in force generation during CME in budding yeast. We quantified the maximum number of molecules of fimbrin and transgelin arriving at CME sites and found that there are around ten times more fimbrin molecules than transgelin molecules on average. We also used an experimentally constrained model of yeast CME to recapitulate the associated actin network assembly and examine the effects of differing numbers of crosslinking proteins on the success of CME. Our modeling results predict that networks with more crosslinking proteins will internalize more effectively, and we showed that in live cells, CME sites with more crosslinking proteins are more likely to internalize against increased resistive forces, corroborating our model prediction. The model also predicts that crosslinking proteins assist in the self-organization of the actin network to optimize the internalization of the plasma membrane driven by actin polymerization, providing a novel role for crosslinking proteins in force generation.

We found that deletion of the gene encoding fimbrin leads to compensatory recruitment of transgelin to sites of endocytosis. To count fimbrin and transgelin molecules, we used a quantitative microscopy technique that allowed us to compare numbers of crosslinking proteins at individual CME sites in various genetic backgrounds. Wild-type yeast recruit about 545 molecules of fimbrin and 40 molecules of transgelin for a total of about 585 crosslinking proteins per CME site. Upon overexpression of transgelin in cells lacking fimbrin, between two and three times the wild-type levels of transgelin are recruited to CME sites. This additional crosslinker abundance partially rescues the endocytic defect caused by the absence of fimbrin.

These results suggest that crosslinker abundance at CME sites is an important factor in determining the success of the actin network in internalizing the vesicle.

Our hypotonic shock experiments indicate that crosslinking proteins are an important force-generating factor for the response to high turgor pressure. Hypotonic shock has been used to increase turgor pressure in studies of endocytosis in yeast in multiple previous studies (6, 26, 28, 29). Deletion of the *FPS1* gene encoding a glycerol transporter leads to a sustained state of increased turgor pressure on the time scale of five to 15 minutes, which causes an endocytic defect (25). In this study, we used hypotonic shock to show that endocytic sites with more crosslinking proteins have a higher likelihood of overcoming the increased energy barrier and internalizing a vesicle, indicating that additional crosslinking proteins generate an actin network capable of generating more force for internalization. A similar example of dosage response in endocytosis has been observed for the type-I myosins, where strains that recruit more myosins to CME sites display increased inward movement speeds during invagination, suggesting that these myosins control the rate of membrane invagination (30). Dosage effects of various other components of the CME pathway may provide novel insights into their mechanisms.

We used mathematical modeling of endocytosis to test the importance and mechanisms of crosslinking proteins in generating actin networks that generate increased forces for internalization. In the model presented in this study, simulated networks that internalize at least 60 nm and reach the snap-through transition point for internalization have more than 2000 total crosslinking proteins bound in the network.

This number is significantly higher than the maximum number of crosslinking proteins measured at sites of CME *in vivo* (∼585). Simulated networks that recruit roughly the physiological number of crosslinking proteins fail to internalize to 60 nm (Fig. 2E). This result suggests that the model is not fully capable of recapitulating the forces generated by the actin network *in vivo* and, therefore, is still missing parts required to generate sufficient force for internalization. The budding yeast type-I myosins are required for CME and generate power through motor domain activity (7, 8), so including myosins in the model could increase the internalization force generated by the actin network, in tandem with actin polymerization and crosslinking proteins. Additionally, internalization is maximized for networks that accumulate around 1500 double-bound crosslinking proteins and decreases slightly for networks with more than that number, suggesting that there is an optimal number of crosslinking proteins for force generation.

Investigation of the nature of actin networks oversaturated with crosslinking proteins both *in silico* and *in vivo* could reveal additional information about their role in CME.

The model presented in this study also predicts that crosslinking proteins assist in self-organization of actin filaments such that growing filament ends are concentrated at the membrane where they are best situated to generate force for internalization. This type of self-organization of actin networks mediated by interactions between actin-associated proteins and actin filaments has been seen in several other systems. Type-II myosin motor activity is needed to coalesce randomly oriented actin filaments at formin nodes into a contractile ring during cytokinesis in fission yeast (31, 32) and nucleation of actin polymerization by ActA from Listeria is sufficient to recruit the necessary downstream actin-binding proteins for assembly of force-producing actin tails (33). The crosslinking protein-directed actin network self-organization observed in this study is maintained even under a high-force regime, suggesting that it may be a critical component of an adaptive response to the varied environmental conditions that budding yeast encounter in the wild. Through the combination of live-cell experiments and mathematical modeling, we uncovered a novel role for actin crosslinking proteins in actin network force generation and organization. Given the high conservation of the proteins involved, we expect that the principles uncovered here will apply broadly to actin networks involved in countless other force-generating processes in a panoply of organisms.

## Methods

### Yeast strains

All budding yeast strains were grown in standard rich media (YPD) at 25°C since *sac6Δ* strains are temperature sensitive. The strains used in this study were derived from the wild-type diploid strain DDY1102 using standard techniques and are listed in Table S1. *SAC6* and *SAC6-GFP* duplication strains were generated following the technique described in Huber et al. (2014) (34). Deletion of *SCP1* was constructed as described previously (35). The *GAL1* promoter was inserted upstream of *SCP1*, disrupting the native promoter, as described previously (35). The resulting strains were verified by DNA sequencing.

### Transgelin Overexpression

Transgelin (Scp1) was overexpressed using the method described in McIsaac et al. (2011) (36). Cells were grown to early log phase in imaging media (synthetic minimal media supplemented with adenine, l-histidine, l-leucine, l-lysine, l-methionine, uracil, and 2% glucose) at 25°C and β-estradiol (Sigma Aldrich) was added to a final concentration of 50 nM to induce overexpression. Cells were then grown for 20 hours and diluted to OD600 = 0.25 in imaging media with 50 nM β-estradiol. Cells were grown for another four hours to mid-log phase before imaging.

### Live-cell imaging

Cells were grown to mid-log phase in imaging media at 25°C and adhered to coverslips coated with 0.2 mg/mL concanavalin A. For quantitative microscopy experiments, reference cells were also grown to mid-log phase in imaging media at 25°C, incubated with FM4-64 (10 μg/mL, Molecular Probes), and adhered to the same coverslips as the reference strain. For hypotonic shock experiments, cells were grown to mid-log phase in imaging media containing 1M sorbitol at 25°C, adhered to coverslips with 0.2 mg/mL concanavalin A, then washed once into imaging media containing 0.25M sorbitol five min before imaging.

Quantitative fluorescence microscopy was performed using a Nikon Ti2-E inverted microscope with an Oko Labs environmental chamber pre-warmed to 25°C. GFP and FM4-64 were excited by a 488-nm laser and mRFP and mCherry were excited by a 561-nm laser using highly inclined and laminated optical sheet (HILO) microscopy to illuminate the medial focal plane of the sample using a LUNF 4-line laser launch (Nikon Instruments, Melville, NY) and an iLas2 TIRF/FRAP module (Gataca Systems, Massy, France). A FITC emission filter was used to filter GFP and a Cy3 emission filter was used to filter mRFP, mCherry, and FM4-64. Movies were acquired using the Nikon NIS Elements software with a Nikon 60X CFI Apo TIRF objective (NA 1.49) and an Orca Fusion Gen III sCMOS camera (Hamamatsu, Hamamatsu City, Japan) at 1 X magnification. A single FM4-64 image was acquired using a 500 ms exposure time and single-channel and dual-channel movies were collected by acquiring images with 500 ms exposure times for 1 min.

Microscopy of Sla1-GFP and Abp1-mRFP was performed using a Zeiss LSM900 with Airyscan 2.0 detection and a stage pre-warmed to 25°C. Two color images of the medial focal plane were acquired using a Zeiss 60X objective (NA 1.4) by Zen Blue software in frame sequential acquisition mode, 5.0x zoom, and maximum laser scan speed at 1.7 fps. GFP and mRFP were imaged with 3% laser power, 850V gain, and 1.0 digital gain. Laser lines were 488 nm for GFP and 561 nm for RFP. Post-acquisition, images were processed in Zen Blue with standard airyscan processing using auto filtering.

### Image Analysis

All image analysis was performed using Fiji software (National Institutes of Health). All movies were subjected to background subtraction and photobleaching correction (5). A median filter was used to quantify GFP-tagged fimbrin and transgelin molecule numbers to subtract cytosolic background from patches, as described previously in Picco and Kaksonen (2017) (37). 2D particle tracking software from the MosaicSuite (38) was used to track fimbrin, transgelin, and Abp1 protein dynamics and trajectories were selected by visual inspection. Trajectories with lifetimes of less than two seconds, or that could not be clearly resolved from another patch, were excluded from our analysis. Maximum fluorescence intensities and fimbrin and transgelin patch lifetimes were extracted from the trajectories using custom code in Python (3.7) with Jupyter Notebook (Project Jupyter). This code is available at the following website: https://github.com/DrubinBarnes/YeastTrackAnalysis. The ratio between fluorescence intensities in the reference and query cells was used to determine molecule numbers as described previously (22).

For quantification of patch internalization and Sla1-GFP and Abp1-mRFP lifetimes, radial kymographs were generated from cells and chosen at random to be measured. The patch was judged as “internalized” if it moved at least 200 nm toward the cell interior. The patch was judged as “failed” if it did not move toward the cell interior, moved fewer than 200 nm toward the cell interior, or moved toward the cell interior before returning to its original position.

### Mathematical modeling in Cytosim

#### Cytosim model

We modified a previously published Brownian dynamics model of budding yeast CME (9) in Cytosim (39). The simulation is bounded by a cylindrical volume centered around an active patch in which all objects diffuse and, upon collision, may associate stochastically based on programmed parameters determined biochemically. The model consists of actin filaments nucleated by Arp2/3 complexes at the plasma membrane and connected by crosslinking proteins. The 3D actin network is bound to a movable object representing the invagination, and actin filaments produce forces on the membrane to internalize the object against a spring-like resistive force. The resistive force at a given internalized distance is described by the equation: *F*(*L*) = *F*_0_ + *k*_*n*_*L*, where F0 is the initial force barrier to internalize the vesicle, kπ is the spring constant, and L is the distance between the end of the vesicle and the membrane surface. In modification of the original model, the invagination object was extended to 120 nm, and the rate of Arp2/3 addition remained constant from three to ten seconds of simulation time. To measure the effect of differing numbers of crosslinking proteins, we varied the number of crosslinking proteins available in the cylindrical volume at the start of the simulation from 0 to 6000. To measure the effect of increasing turgor pressure, we varied the spring constant, kπ, on the invagination object from 13,333 pN/nm to 30,000 pN/nm.

#### Running simulations

Custom scripts in bash were used to run parallel simulations on a high-performance computing server.

#### Analysis of simulations

Custom code in Python (3.7) with Jupyter Notebook (Project Jupyter) was used to report, read, analyze, and plot the simulations obtained from Cytosim. X, Y = 0 is defined as the center of the pit, and Z = 0 is defined as the membrane. The potential polymerization energy of the network at a given time point was defined as the sum of the potential polymerization energy of all growing actin filaments within a distance of 1.375 nm from the membrane. The potential polymerization energy of an individual actin filament was defined as 9 pN * 2.75 nm * sinθ where θ is the angle of incidence with the membrane and θ = 0 is defined as parallel with the membrane. This code is available at the following website: https://github.com/DrubinBarnes/Hill_Cai_Carver_Drubin_2024_Manuscript.

## Acknowledgements

We thank Serge Dmitrieff, Francois Nedelec, and Matt Akamatsu for providing code and configurations for running and analyzing simulations using Cytosim. We thank Padmini Rangamani, Yidi Sun, Jonathan Wong, and Julia Torvi for advice and support. In addition, we thank the UC Berkeley High-Performance Computing cluster for training and server space for parallel simulations. We thank Max Ferrin, Ross Pedersen, Thomas Pollard, Yidi Sun, and our anonymous reviewers for their critical reading of the manuscript. This research was conducted with US Government support, under National Institutes of Health Grant R35GM118149 to D.G.D.

---

**Fig. S1.**
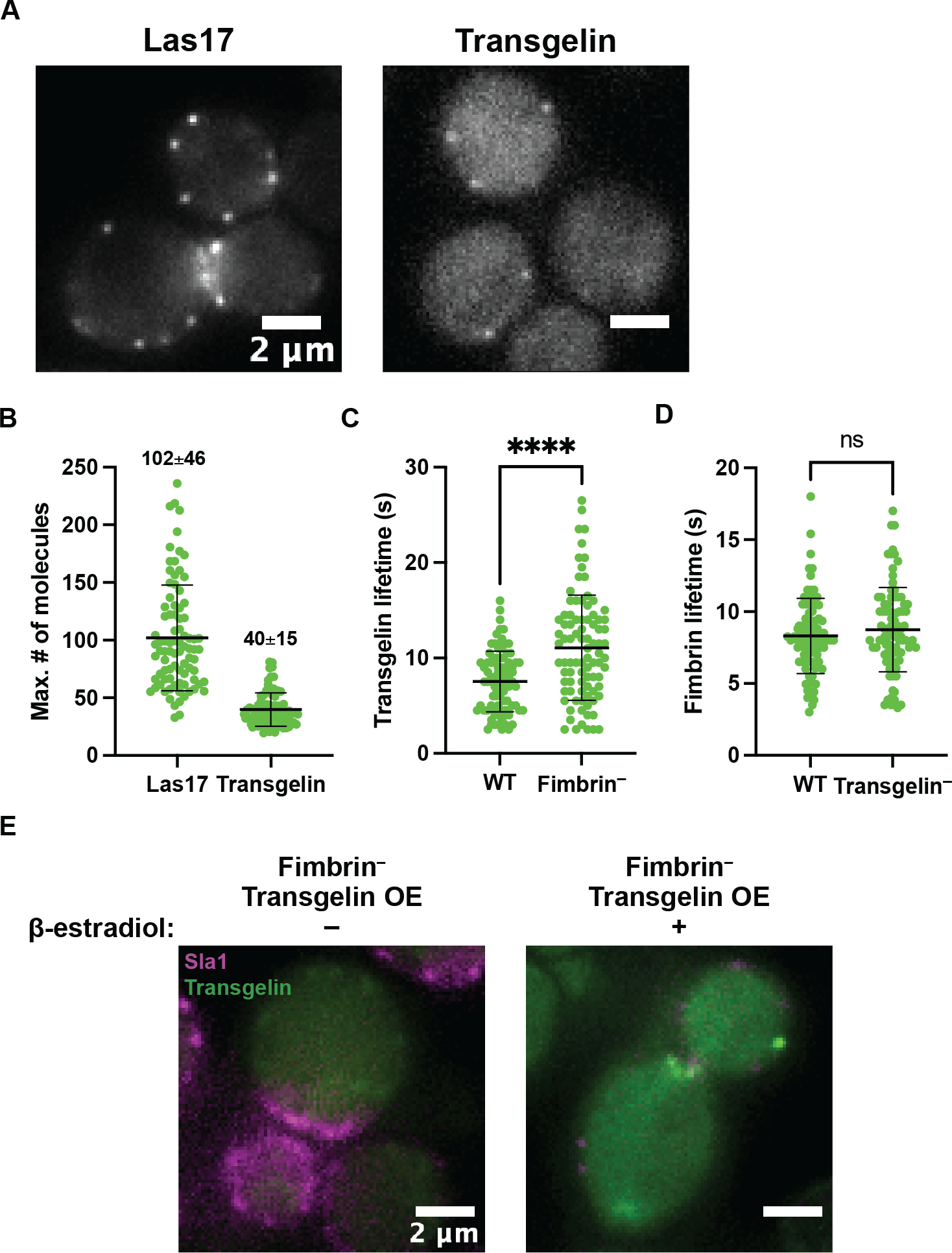
Quantification of maximum number of transgelin molecules arriving at CME sites. (A). Quantitative microscopy of GFP-tagged Las17 (left) and transgelin (right). (B). Quantification of maximum numbers of Las17 molecules and transgelin molecules arriving at CME sites. (C), (D). Lifetimes for transgelin (C) and fimbrin (D) patches at endocytic sites for wild-type and strains lacking fimbrin (fimbrin^−^) or transgelin (transgelin^−^), respectively (n=90 sites from ≥10 cells per strain). Error bars show standard deviation. (E). Representative images of mCherry-tagged Sla1 and GFP-tagged transgelin in a fimbrin^−^ transgelin overexpression strain (transgelin OE), without induction by β-estradiol (left), and 24 hours post-induction with β-estradiol (right).

**Fig. S2.**
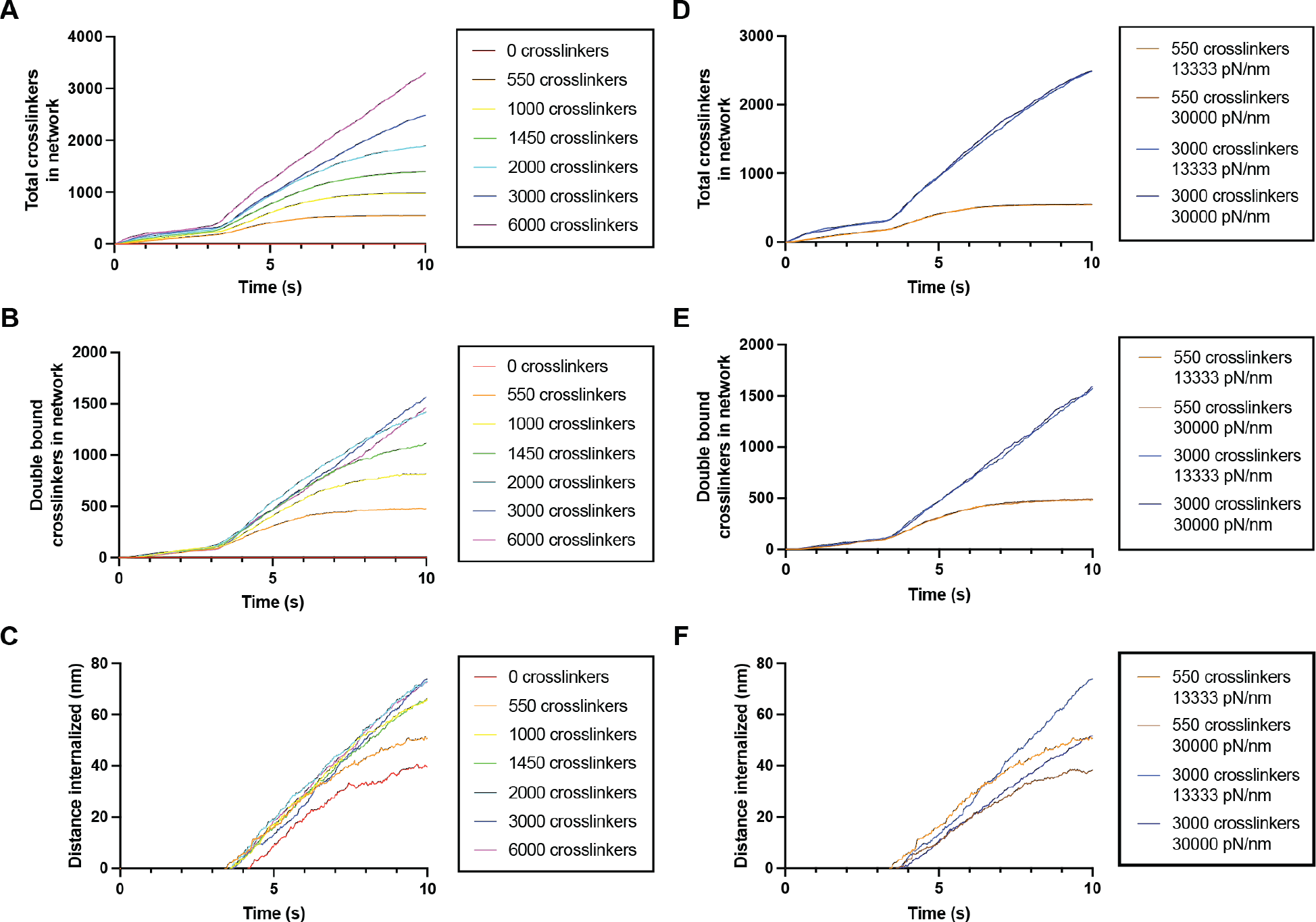
Total and double-bound crosslinkers and vesicle internalization for simulations with various pools of available crosslinking proteins. (A-C) Total number of crosslinking proteins with at least one actin-binding site bound to an actin filament (A), double-bound crosslinkers (B), and distance of vesicle internalization (C) over the duration of simulations with a range of pools of available crosslinking protein numbers with 13333 pN/nm required to internalize the vesicle. N=5 simulations. (D-F) Total number of crosslinking proteins with at least one actin-binding site bound to an actin filament (D), double-bound crosslinkers (E), and distance of vesicle internalization (F) over the duration of simulations with pools of 550 or 3000 available crosslinking proteins and 13333 pN/nm or 30000 pN/nm required to internalize the vesicle. N=5 simulations.

**Fig. S3.**
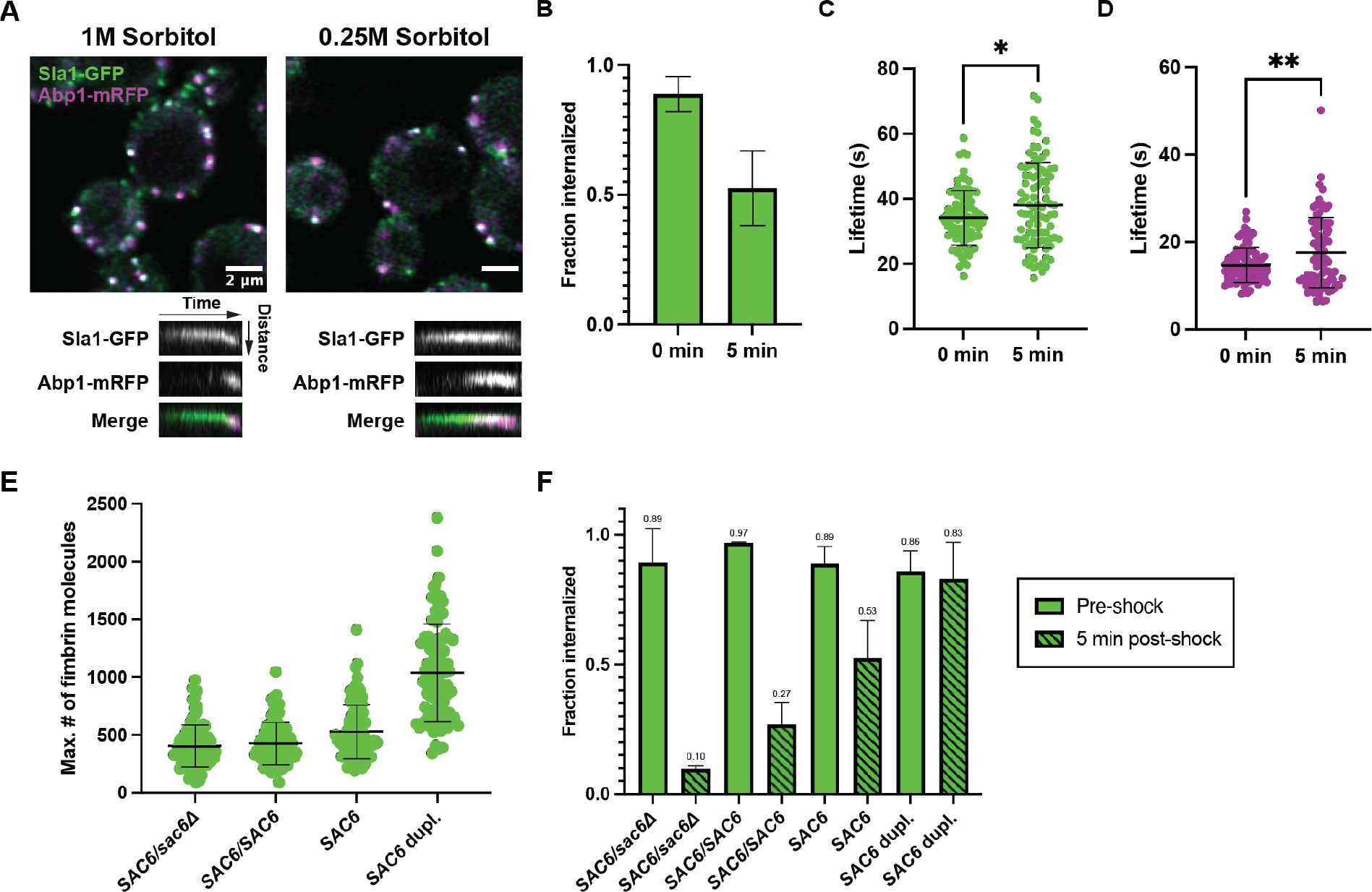
Hypotonic shock stalls endocytosis. Experiment performed as in Fig. 3. (A) WT cells grown in media containing 1M sorbitol (left) or five minutes after shifting to media containing 0.25M sorbitol (right). Representative kymographs are shown for each condition. (B) Fraction of CME sites internalized before and after hypotonic shock. Sites where Sla1 and Abp1 hook into the cell were considered internalized (n=90 sites from ≥10 cells). C, D. Lifetimes of GFP-tagged Sla1 (C) and mRFP-tagged Abp1 (D) at endocytic sites (n=90 sites from ≥10 cells). (E) Quantification of the maximum number of fimbrin molecules arriving at endocytic sites five minutes after hypotonic shock for strains with different copy numbers of the gene encoding fimbrin (n=90 sites from ≥10 cells). (F) Fraction of sites internalized before and after hypotonic shock for strains with different copy numbers of the gene encoding fimbrin (n=3 technical replicates). Error bars show standard deviation.

**Fig. S4.**
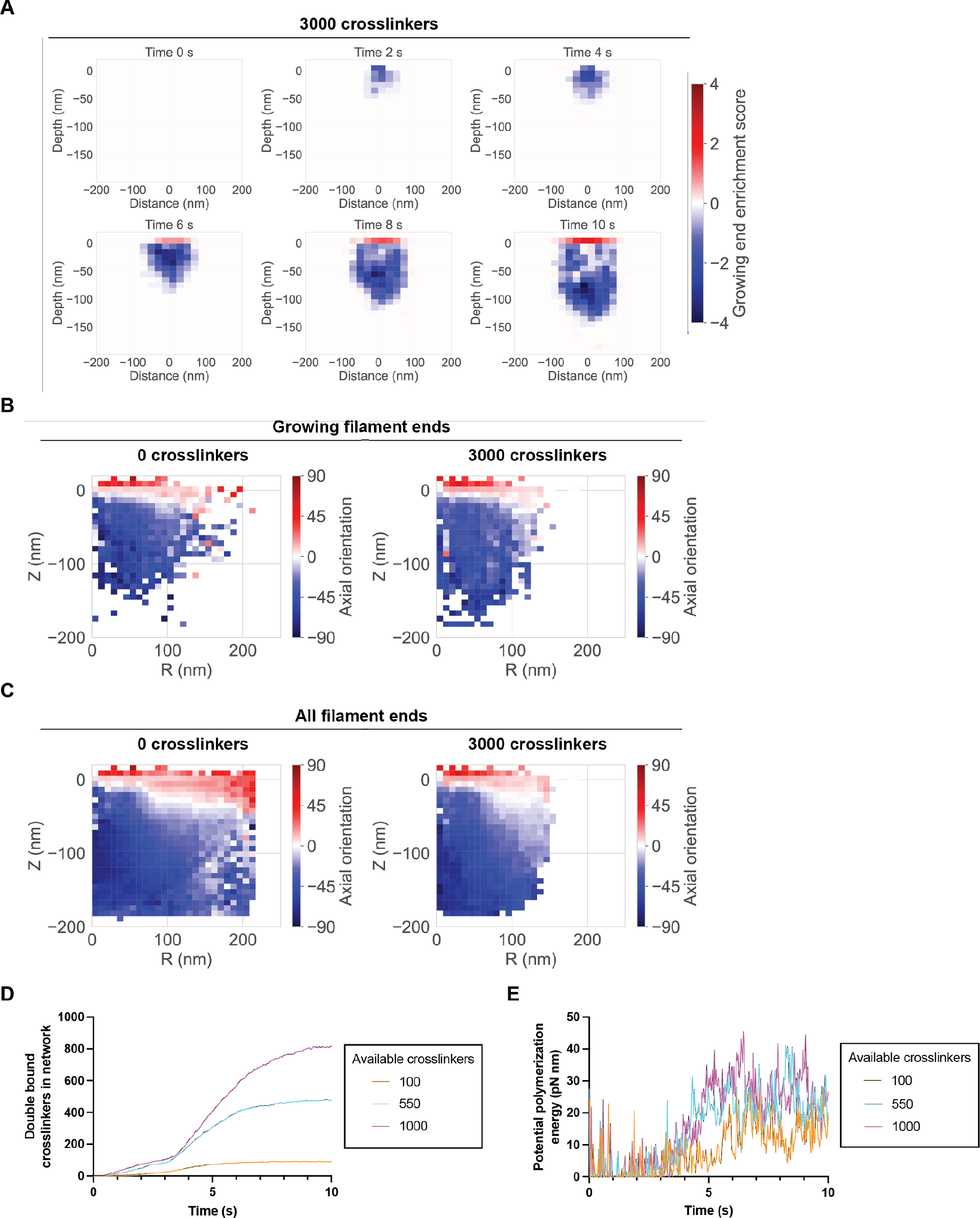
Filament enrichment, axial orientations, and low crosslinking protein simulations. (A) Heat map of enrichment scores for growing actin filament ends relative to capped ends with a pool of 3000 available crosslinking proteins in two-second increments throughout the length of the simulation. N= 5 simulations. (B) Axial orientation of growing filament ends measured radially from the center of the invagination for simulations with pools of 0 or 3000 available crosslinking proteins. N=5 simulations. (C) Axial orientation of both growing and capped filament ends measured radially from the center of the vesicle for simulations with pools of 0 or 3000 available crosslinking proteins. N=5 simulations. (D) Double-bound crosslinkers over the duration of simulations in the low available crosslinking protein regime. N=5 simulations. (E) Average potential polymerization energy of the actin network over the duration of simulations in the low available crosslinking protein regime. N=5 simulations.

**Table S1.** **Strains used in this study**

| Name | Genotype | Source |
| --- | --- | --- |
| DCT575 | *MATa, his3-Δ200, leu2-3, 112, ura3-52, SCP1-GFP::KanMX* | Drubin Lab |
| DDY2736 | *MATa, his3-Δ200, leu2-3, 112, ura3-52, lys2-801, LAS17-GFP::HIS3,* | Drubin Lab |
| LCY8.1 | *MATα, his3-Δ200, leu2-3,112, ura3-52, SCP1-GFP::HIS3, sac6Δ::cgLEU2* | This study |
| LCY20.1 | *MATα, his3-Δ200, leu2-3,112, ura3-52, lys2-801, scp1Δ::cgURA3, SAC6-GFP::HIS3* | This study |
| DDY3624 | *MATα, his3-Δ200, leu2-3,112, ura3-52, SAC6-GFP::HIS3* | Drubin Lab |
| JHY244.1 | *MATα, his3-Δ200, leu2-3,112, ura3-52, SCP1-GFP::HIS3, sac6Δ::cgLEU2, Sla1-mCherry::HIS3* | This study |
| JHY246.1 | *MATα, his3-Δ200, leu2-3,112, ura3-52, SCP1-GFP::HIS3, Sla1-mCherry::HIS3* | This study |
| JHY248.1 | *MATα, his3-Δ200, leu2-3,112, ura3-52, HygR::PGAL1-SCP1-GFP::HIS3, sac6Δ::cgLEU2, Sla1-mCherry::HIS3, GEV::URA3* | This study |
| JHY174.1 | *MATα, his3-Δ200, leu2-3,112, ura3-52, scp1Δ::cgURA3, fps1Δ::natNT2, SLA1-GFP::KanMX6, ABP1-mRFP::HIS3* | This study |
| JHY183.1 | *MATα/MATa, his3-Δ200/-, leu2-3,112/-, ura3-52/-, lys2-801/+, scp1Δ::cgURA3/-, fps1Δ::NatNT2/-, SAC6-GFP::HIS3/sac6Δ::cgLEU2* | This study |
| JHY184.1 | *MATα/MATa, his3-Δ200/-, leu2-3,112/-, ura3-52/-, scp1Δ::cgURA3/-, fps1Δ::NatNT2/-, SAC6-GFP::HIS3/SAC6-GFP::HIS3* | This study |
| JHY182.1 | *MATα, his3-Δ200, leu2-3,112, ura3-52, scp1Δ::cgURA3, fps1Δ::NatNT2, SAC6-GFP::HIS3* | This study |
| JHY218.1 | *MATα, his3-Δ200, leu2-3,112, ura3-52, lys2-801, scp1Δ::cgURA3, fps1Δ::NatNT2, SAC6-GFP::HIS3::cgLEU2::SAC6-GFP::HIS3* | This study |
| JHY178.1 | *MATα/MATa, his3-Δ200/-, leu2-3,112/-, ura3-52/-, lys2-801/+, scp1Δ::cgURA3/-, fps1Δ::natNT2/-, SLA1-GFP::KanMX6/+, ABP1-mRFP::HIS3/+, sac6Δ::cgLEU2/+* | This study |
| JHY179.2 | *MATα/MATa, his3-Δ200/-, leu2-3,112/-, ura3-52/-, scp1Δ::cgURA3/-, fps1Δ::natNT2/-, SLA1-GFP::KanMX6/+, ABP1-mRFP::HIS3/+* | This study |
| JHY176.1 | *MATα, his3-Δ200, leu2-3,112, ura3-52, scp1Δ::cgURA3, fps1Δ::natNT2, SAC6::HygMX4::SAC6, SLA1-GFP::KanMX6, ABP1-mRFP::HIS3* | This study |
| JHY162.1 | *MATα, his3-Δ200, leu2-3,112, ura3-52, lys2-801, scp1Δ::cgURA3, fps1Δ::NatNT2, ABP1-mRFP::HIS3, SAC6-GFP::HIS3* | This study |

